# Design of peptide-based PAC1 antagonists combining molecular dynamics simulations and a biologically relevant cell-based assay

**DOI:** 10.1101/2025.04.16.649181

**Authors:** Wenqin Xu, Abigail M. Keith, Wenjuan Ye, Xin Hu, Noel Southall, Juan J. Marugan, Marc Ferrer, Mark J. Henderson, Patrick M. Sexton, Giuseppe Deganutti, Lee E. Eiden

## Abstract

The PACAP receptor PAC1 is a G_s_-coupled family B1 GPCR for which the highest-affinity endogenous peptide ligands are the pituitary adenylate cyclase-activating peptides PACAP38 and PACAP27, and whose most abundant endogenous ligand is PACAP38. PACAP action at PAC1 is implicated in neuropsychiatric disorders, atherosclerosis, pain chronification, and protection from neurodegeneration and ischemia. As PACAP also interacts with two related receptors, VPAC1 and VPAC2, highly selective ligands, both agonists and antagonists, for PAC1 have been sought. To date, the peptide PACAP(6-38) and polypeptide M65, which is related to maxadilan, a sandfly vasodilator peptide, have been identified as selective for PAC1. Several non-peptide small molecule compounds (SMOLs) have been reported to be specific antagonists at PAC1, albeit their specificities have not been rigorously documented. Here, we present a platform of cellular assays for the screening of biologically relevant antagonists at PAC1 and show that some currently proposed SMOL antagonists do not have activity in this cell reporter assay, while we confirm that PACAP(6-38) and M65 are competitive antagonists. We have used this assay system to explore other peptide antagonists at PAC1, guided by molecular dynamics analysis of the PACAP-PAC1 interaction based on cryo-EM structural models of PAC1 complexed with a number of biologically active ligands. The affinity-trap model for the PAC1-ligand interaction successfully predicts the engagement behavior of PACAP27 and PACAP38 peptide-based PAC1 inhibitors. In particular, C-terminal deletants of PACAP(6-38) that maintain equipotency to PACAP(6-38) allow the shorter sequence to function as a scaffold for further peptide-based antagonist exploration.

## 1. Introduction

PACAP action at its receptor PAC1 has been implicated in a variety of physiological contexts relevant to human health and disease. These include heart disease, where PACAP is proposed to act at PAC1 to attenuate inflammatory atrial fibrosis and atherosclerogenesis [1], neuropsychiatric disorders, in particular depression and post-traumatic stress disorder [2], migraine [3], dry eye syndrome [4], nerve injury-induced allodynia [5], and neuroprotection from tissue damage in stroke and ischemia [6–10]. As implied by its broad actions as a regulatory peptide, therapeutic application of both PAC1 agonists and antagonists can be envisaged. Exploration of the basis of PACAP selectivity at PAC1 is a necessary first step in the development of agonist and antagonist compounds, both for potential therapeutic application and as pharmacological tools to explore where and when, in various disease processes, PACAP might facilitate or attenuate pathophysiology.

Structural analysis of PAC1 has revealed that binding of PACAP has both similarities and differences compared to ligand binding to closely related family B1 receptors, for which high-resolution structures have been obtained [11–13]. Similarities include initial binding events between the C-terminal domain of the peptide with the extracellular domain (ECD) followed by insertion of the N-terminus of the peptide into the core of the receptor to enable activation, but with differences in binding poses in the C-terminus explaining in part the ligand selectivity of each receptor [13]. Empirically, prior to structural analysis, PACAP(6-38), hereafter also referred to as P6-38, was identified as an antagonist [14] and maxadilan as an agonist [15], each with specificity for PAC1, although P6-38 can also antagonize PACAP action at VPAC2 [16]. Deletion of the central portion of maxadilan produces the antagonist M65, consistent with the cryo-EM structure of maxadilan-PAC1 in which maxadilan binds to the receptor via antiparallel helices with the central domain protruding into the receptor core to produce activation of G_s_ [17]. A model has been formulated for PACAP binding based on both pharmacology and structural analysis that is consistent with models for other B1 GPCRs, in which the C-terminus of PACAP engages the ECD and extracellular loop 1 (ECL1) of PAC1, with the N-terminal domain then positioning within the receptor core to effect activation [12, 18, 19]. This ‘affinity trap’ model emphasizes that initial binding to the ECD is relatively well-modeled by NMR and X-ray crystallographic structure determinations based on ECD alone in complex with peptide. An ECD-based model has also been used for *in silico* identification of potential PACAP antagonists [20, 21].

There has been success in the development of small molecule compounds (SMOLs) as agonists for GLP-1 and GCGR [22] and a non-peptide antagonist for CRF1R [23]. However, the yield of SMOL antagonists based on the ECD-ECL1 high-affinity binding site for other family B1 (‘secretin-like’) receptors has been relatively meager, as have those based upon competitive inhibition of PACAP-receptor binding, or simply empirical screening of both competitive and allosteric small-molecule inhibitors in cell-based screens [24–26].

Furthermore, PAC1 antagonist SMOLs have not been widely characterized for off-target effects, nor used widely beyond the laboratories in which they were originally discovered. Likewise, peptide-based agonists and antagonists for PAC1 have not been extensively explored since initial identification of P6-38 and M65, with the exception of the maxadilan-conjugated MAXCAP (US patent #12049486B2), and the PACAP-derived recombinant peptide MPAPO [27]. This is perhaps in part due to early structure-activity analysis of PACAP38 and 27 deletants prior to detailed structural knowledge of the peptide-receptor complex structures, for example the finding that sequential removal of amino acids from the C-terminus of P6-38 causes rapidly diminishing binding to PAC1 and VPAC2 [28]. These studies, moreover, were performed using cell-free receptor binding assays in which the binding properties of PACAP to its receptor that occur in a cellular context may not all be fully captured.

We decided to look carefully at existing SMOLs reported as PACAP antagonists, and to ‘back-validate’ a cellular assay based on activation by maxadilan, PACAP38 (P38), and PACAP27 (P27), as well as the antagonism by P6-38 and M65, the most frequently reproduced and selective antagonists for *in vivo* PACAP signaling yet reported. The development of a biologically relevant cell-based assay for PAC1 activation, and its deployment in the testing of structural predictions for PACAP binding to PAC1 based on molecular dynamics simulations analysis, is reported here.

## 2. Materials and Methods

### 2.1 Drugs

Peptides including vasoactive intestinal peptide (VIP), pituitary adenylate cyclase-activating polypeptide (PACAP) 1-38 (P38), 1-27 (P27), 6-38 (P6-38), 6-27 (P6-27), and a series of C-terminal deletants of P6-38 (6-36, 6-32, 6-30, 6-29 and 6-28), were synthesized and purchased from AnaSpec (Fremont, CA). The following P6-27 analogues, featuring a lactam bridge between Glu16 and Lys20, and P6-38 analogues, containing a lactam bridge between Glu25 and Lys29, were also synthesized and obtained from AnaSpec: 6-27Lb: (c[E16,K20])P6-27; P6-27Lbx1 with a Cha substitution at position 23: (c[E16,K20], Cha23)P6-27; P6-27Lbx2 with a Cha substitution at position 27: (c[E16,K20], Cha27)P6-27; P6-38Lb: (c[E25,K29])P6-38; P6-38Lbx1 with a Cha substitution at position 23: (c[E25,K29], Cha23)P6-38.

Maxadilan and M65 were purchased from Bachem (Torrance, CA), and forskolin was obtained from Tocris (Minneapolis, MN).

Published SMOLs identified as PACAP antagonists [20, 21, 25, 26] were obtained from various commercial sources. Beebe-1 and Beebe-2 were purchased from Enamine (Monmouth Junction, NJ). PA-8 was acquired from two separate vendors: Tocris (referred to as PA-8-t in this study) and MedChemExpress (Monmouth Junction, NJ) (referred to as PA-8-m). PA-9 was obtained from the NCATS in-house compound library. PA-9 analog 3d (also named as PA-915) was obtained from MedChemExpress, while PA-8 analog 2o was purchased from Enamine. Bay2686013 was also obtained from Enamine.

Drug stocks were prepared according to manufacturers’ instructions, either in assay medium or DMSO. A vehicle control with the same DMSO concentration as the treatments was included whenever appropriate.

### 2.2 Cell Culture

Cell culture reagents were purchased from ThermoFisher Scientific (Waltham, MA) unless otherwise noted. Neuroscreen-1 (NS-1) cells (Cellomics, Halethorpe, MD) were grown on flasks or plates coated with Collagen I (rat tail) as described previously [29]. NS-1 cells were cultured in RPMI-1640 media supplemented with 10% horse serum and 5% heat-inactivated Fetal Bovine Serum (Hyclone, Logan, UT). HEK293 and SH-SY5Y cells were purchased from ATCC (Gaithersburg, MD) and cultured with DMEM medium with 10% FBS. All cell culture medium were supplemented with 2 mM glutamine, 100 U/ml penicillin, and 100 μg/ml streptomycin.

### 2.3 Establishment of stable cell lines

A single clone of the SH-SY5Y cell line stably expressing a cAMP biosensor (SY5Y_CBS) was generated as previously described for HEK293_CBS and NS-1_CBS cells [30], using the retroviral vector pLHCX containing the cAMP biosensor and a hygromycin resistance gene. A HEK293_CBS_hPAC1 cell clone was generated by transducing HEK293_CBS cells with the lentiviral vector pLX304 (GeneCopoeia, Rockville, MD) encoding the human PAC1hop (PAC1; GenBank accession no. BC143679) and a blasticidin resistance gene . Following transduction, cells were selected with the appropriate antibiotics (hygromycin or blasticidin) for 2 weeks. Single-cell clones were subsequently isolated by limiting dilution and assessed for PACAP-induced cAMP production using concentration-response assays.

### 2.4 Neuritogenesis assay

NS-1 cells were seeded in 6-well plates at a density of 50,000 cells per well and incubated overnight. Cells were pre-treated with PACAP antagonists, including individual P6-38 deletants, for 30 minutes prior to the addition of 100 nM PACAP. The compound solvent was used as a vehicle control. After 48 hours, ten micrographs per well were automatically acquired at 20× magnification using the Jobs module of the NIS-Elements acquisition software (version 4.51.01, Nikon, Edgewood, NY) on a Nikon ECLIPSE Ti microscope. Neurite length and cell number were quantified using NIS-Elements AR Analysis (version 5.42.03) with the AI Image Processing module. Briefly, the *segment objects.ai* tool was trained to generate a binary layer that identified cell bodies, which were then automatically counted to determine the cell number per image. Separately, the *segment.ai* tool was trained to produce a binary layer that traced neurites, allowing measurement of total neurite length per image. Prior to quantification, all binary layers were manually reviewed and edited to correct any AI-generated errors. The average neurite length per cell was calculated for each image and then averaged across the ten images to determine the average neurite length per cell for each well. The reported values represent the mean ± SEM from three independent experiments performed on different days, with triplicate wells for each treatment in an individual experiment (n = 3).

### 2.5 Cyclic AMP measurement

The cyclic AMP biosensor (CBS) assay was performed using Gibco CO₂-independent medium (ThermoFisher, catalog #18045-088). HEK293_CBS_hPAC1 or SY5Y_CBS cells were seeded in 96-well black/clear-bottom plates (MidSci, Fenton, MO) at a density of 6,000 cells per well and incubated overnight. The following day, the medium in each well was aspirated and replaced with fresh medium containing 1.5 mg/mL D-luciferin potassium salt substrate (Gold Biotechnology, St. Louis, MO) and 500 µM isobutyl-1-methylxanthine (IBMX), followed by incubation at 37 °C for 1 hour. Cells were then pre-treated with antagonists for 10 minutes before the addition of agonists (PACAP or forskolin). Plates were incubated at 37 °C for 1 hour, then brought to room temperature and equilibrated for 30 minutes. Luminescence was measured using a GloMax plate reader (Promega, Madison, WI).

For the high-throughput CBS screening assays, cells were plated at 2,000 (HEK293_CBS_hPAC1) or 3,000 (SY5Y_CBS) cells per well in 4 mL of Gibco CO_2_-independent medium, in white 1,536-well tissue culture treated plates. After overnight incubation at 37 °C and 5% CO_2_, 4 mL of luciferin/IBMX solution was added to the wells (final concentrations of 500 µM each) and plates were incubated at 37 °C for 1 hour. SMOLs (22.5 nL) were transferred to the assay plate, using a pintool device (HEK293 screen) or an ECHO acoustic liquid handler (Beckman Coulter, Brea, CA, SY5Y screen), to final concentrations (4 concentrations per compound) between 0.5 and 30 µM and cells were incubated at 37 °C for 1 hour (HEK293 screen) or 10 minutes (SY5Y screen). Agonist peptide (P27 or P38), was then added to each well by dispensing 1 mL of peptide diluted in medium (HEK293, final concentration 2 nM) or by acoustic dispenser (SY5Y, final concentration 0.2 nM). For forskolin counterscreening in SY5Y cells, forskolin instead of peptide was dispensed to the plate at a final concentration of 5 µM.

The NCATS library compound collection (156,059 compounds) [31], used for high-throughput screening with HEK293_CBS_hPAC1 cells, included the NCATS Pharmaceutical Collection (NPC, approved drugs), at four concentrations. Active hits with >50% P38 inhibition were selected for confirmation assay at seven concentrations, with forskolin stimulation as a counter assay. Hits were selected based on reproducible inhibition of >50% P38 activity at the highest concentration tested (30 µM), yielding 192 compounds for further testing. Selected hits were followed up and tested in SY5Y_CBS cells. In addition, a total of 2,816 compounds from the NCATS HEAL library, which includes drugs, probes, and tool compounds that act on published pain and addiction-relevant targets [32] were screened in SY5Y_CBS cells at three concentrations. Direct cAMP determinations were made using the DetectX® Direct Cyclic AMP Enzyme Immunoassay Kit (Arbor Assays, Ann Arbor, MI). SY5Y_CBS cells (200,000 cells per well) or HEK293_CBS cells (10,000 cells per well) were seeded into 96-well plates one day prior to the assay. These cell densities were optimized to ensure agonist responses fell within the linear range of the cAMP standard curve (0.16-10 pmol/mL cAMP). On the day of the assay, the medium was removed and replaced with fresh medium containing 500 µM IBMX. After a 20-minute incubation at 37 °C, antagonists were added and incubated for 10 minutes at 37 °C, followed by the addition of agonists for 30 minutes at 37 °C. Following the final incubation, the medium was aspirated, and cells were lysed with 200 µL of lysis buffer (provided in the kit) for 10 minutes at room temperature. Samples were then snap-frozen and stored at -80 °C overnight and thawed the next day to ensure complete cell lysis. The assay was completed by sequentially adding the cAMP peroxidase conjugate, anti-cAMP sheep antibody, substrate, and stop solution, according to the manufacturer’s instructions. Optical density was measured at 450 nm using a microplate spectrophotometer (Molecular Devices, San Jose, CA). Standard curves and data analysis were performed using GraphPad Prism with a sigmoidal four-parameter nonlinear regression model. Experiments conducted on different days with independently plated and assayed cells were considered biological replicates. Each experiment included three technical replicates.

### 2.6 Data Analysis

GraphPad Prism ver. 9.0 or 10.0 (GraphPad Software Inc, Boston, MA) was used for data analysis. Concentration response curves were created using a four parameter logistic regression model. EC_80_ and IC_50_ parameters were obtained using non-linear regression curve fitting tools. EC_80_ was calculated as the effective concentration of the agonist required to produce 80% of the maximal response. IC_50_ was calculated as the concentration of the antagonist that produced 50% of the response induced by an agonist (either PACAP or forskolin) in the absence of the antagonist. Seven or eight different concentrations and three technical replicates per concentration were used for curve fitting. All data are presented as mean ± standard error of the mean (SEM). The tables for EC_80_ or IC_50_ calculation were derived from three independent assays, performed with technical triplicates, for n = 3.

### 2.7 Molecular Modeling of P6-38 and Derivatives

The structure of the P38:PAC1 complex was modeled into the published EM map EMD-20278 [33]. The ECD was rigid body fitted from the X-ray structure of the ECD of PAC1 (PDB: 3N94). Complexes between PAC1 and P6-38 or its derivatives P6-38Lb, P6-38Lbx, and P6-30 were modeled from the P38:PAC1 complex after removal of the G_s_. P6-38 was modeled by removing P38 residues His1-Ile5. P6-38Lb was modeled by removing P38 residues His1-Ile5, mutating Ala25 to a Glu and reorienting both Glu25 and Lys29 side chains to a favorable position for an amidic bond (later introduced as a CHARMM patch; see below). P6-38Lbx1 was modeled like P6-38Lb with an L-cyclohexylalanine (Cha) residue in place of Leu23. P6-30 was modeled by removing P38 residues His1-Ile5 and Tyr31-Lys38. P6-38Lb Ala25Glu substitution was performed with Chimera 1.14 [34]. P6-38Lb side-chain reorientation and P6-38Lbx1 Cha side-chain building was performed manually with Chimera 1.14.

### 2.8 Complexes Preparation for Molecular Dynamics Simulations

PAC1:P6-27, PAC1:P6-27Lb, PAC1:P6-27Lbx1, PAC1:P6-27Lbx2, PAC1:P6-38, PAC1:P6-38Lb, PAC1:P6-38Lbx1, and PAC1:P6-30 complexes were oriented according to the PAC1nR coordinates retrieved from the OPM database [35] before undergoing preparation with the CHARMM36 force field [36]. Each complex was prepared for simulations using in-house scripts based on HTMD2.3.2 [37] and VMD1.9.4 [38] frameworks. This multistep procedure performs the preliminary hydrogen atoms addition employing the pdb2pqr [39] and PROPKA3 [40] software combination through the systemPrepare HTMD implementation [41], considering a simulated pH of 7.4. Following this, the receptors were then embedded in a rectangular 1- palmitoyl-2-oleyl-sn-glycerol-3-phosphocholine (POPC) bilayer (previously built by using the VMD Membrane Builder plugin 1.1 at http://www.ks.uiuc.edu/Research/vmd/plugins/membrane/) considering the coordinates retrieved from the OPM database to gain the correct orientation within the membrane, while removing the lipid molecules overlapping the receptor TMD bundle. TIP3P water molecules [42] were added to the simulation box using the VMD Solvate plugin 1.5 (VMD Solvate plugin, Version 1.5; http://www.ks.uiuc.edu/Research/vmd/plugins/solvate/). Finally, the overall charge neutrality was reached by adding Na^+^/Cl^−^ counter ions (final ionic concentration of 0.150 M) using the VMD Autoionize plugin 1.3 (Autoionize Plugin, Version 1.3; http://www.ks.uiuc.edu/Research/vmd/plugins/autoionize/).

### 2.9 Equilibration and Molecular Dynamics Production Simulations

For both systems, the equilibration and production simulations were computed using the ACEMD3 [43] MD engine. Systems (Table MD1) were equilibrated in isothermal-isobaric conditions (NPT) using the Monte Carlo barostat [44] with a target pressure of 1 atm, the Langevin thermostat [45] with a target temperature of 310 K, along with a low damping factor of 1 ps^−1^ and an integration time step of 2 fs. Clashes between protein and lipid atoms were reduced through 1,500 conjugate-gradient minimization steps, followed by a 4 ns long MD simulation with a linearly-released positional constraint of 1 kcal mol^−1^ Å^−2^ on protein and lipid phosphorus atoms. Subsequently, 80 ns of MD simulation were performed, constraining only the protein atoms. Lastly, positional constraints were applied only to the protein backbone alpha carbons for a further 20 ns. Productive trajectories (in triplicate, 250 ns for each replica) were computed with an integration time step of 4 fs in the canonical ensemble (NVT). The temperature was set at 310 K, using a thermostat damping of 0.1 ps^-1^ and the M-SHAKE algorithm [46] to constrain the bond lengths involving hydrogen atoms. The cut-off distance for electrostatic interactions was set at 9 Å, with a switching function applied beyond 7.5 Å. Long-range Coulomb interactions were handled using the particle mesh Ewald summation method (PME) [47] by setting the mesh spacing to 1.0 Å. Frames were saved every 100 ps. The composition of the simulated systems is detailed in Table MD1.

### 2.10 Mixed Molecular Dynamics Simulations of PA-8 and PA-9

PAC1 in the unliganded, P38-free form after remotion of G_s_ was oriented according to the PAC1n coordinates retrieved from the OPM database, prepared as reported above in Section 2.8 and equilibrated as reported in Section 2.9. The last frame of the equilibration was extracted, and twenty PA-8 and PA-9 molecules, in either R or S isomeric form, were inserted in the resulting apo PAC1 simulation box using PACKMOL [48] (tolerance = 2.0, nloop = 10, radius 1.5). PA-8 and PA-9 molecules overlapping the receptor or the POPC were removed, leaving six or seven SMOLs per system. This generated four new simulation boxes (Table MD1). VPAC1, also modeled in the apo form, was retrieved from PDB 8E3Y [19] and prepared with nine (S)-PA-9 molecules as reported above for PAC1R. The four PAC1:SMOLs and the VPAC1:(S)-PA-9 system were further equilibrated for one nanosecond in nPT without restraints before seeding productive simulation replicas (Table MD1).

### 2.11 MD analysis

Root mean square deviations (RMSD) were computed with VMD1.9.4. Interatomic contacts and hydrogen bonds were detected using the GetContacts scripts tool (https://getcontacts.github.io), setting a hydrogen bond donor-acceptor distance of 3.3 Å and an angle value of 120° as geometrical cut-offs. Contacts and hydrogen bond persistency are quantified as the percentage of frames (over all the frames obtained by merging the different replicas) in which protein residues formed contacts or hydrogen bonds with the ligand. Volumetric maps for SMOLs were computed on the merged trajectories using the Volmap VMD plugin (version 1.1; https://www.ks.uiuc.edu/Research/vmd/plugins/volmapgui/). The MMPBSA.py script [49] from the AmberTools20 was used to compute the VIP binding energy as molecular mechanics energies combined with the generalized Born and surface area continuum solvation (MM/GBSA) method, after transforming the CHARMM psf topology files to an Amber prmtop format using ParmEd (documentation at http://parmed.github.io/ParmEd/html/index.html).

## 3. Results

### 3.1. Binding of PAC1 with PACAP and its antagonists

Figure 1 depicts the structure of PAC1, formed by a seven-helix transmembrane domain (TMD) and an ECD. Binding regions for agonists P27, P38, and maxadilan, as characterized by cryo-EM, span both ECD and TMD [17, 19]. Peptide-based antagonists include an N-terminally truncated version of P38, P6-38, an N-terminally truncated version of P27, P6-27, and M65, a version of maxadilan in which the mid-region of the peptide, which docks to the activation pocket of PAC1, is deleted and the C- and N-terminal helices binding to PAC1 are covalently joined [17, 50]. The N-terminally truncated peptide antagonists are predicted to bind in a fashion similar to agonists. Indeeed, the activity of P6-27 or P6-38 as antagonists is consistent with a model of PACAP binding to PAC1 in which the peptide N-terminus (residues 1-5) is required for receptor activation and G_s_ protein signaling, while the C-terminus is involved in initial binding (affinity-trapping) to PAC1. Previous work predicted SMOLs to bind to PAC1 and inhibit PACAP binding and receptor activation, in both ECD and TMD regions [20].

**Figure 1.**
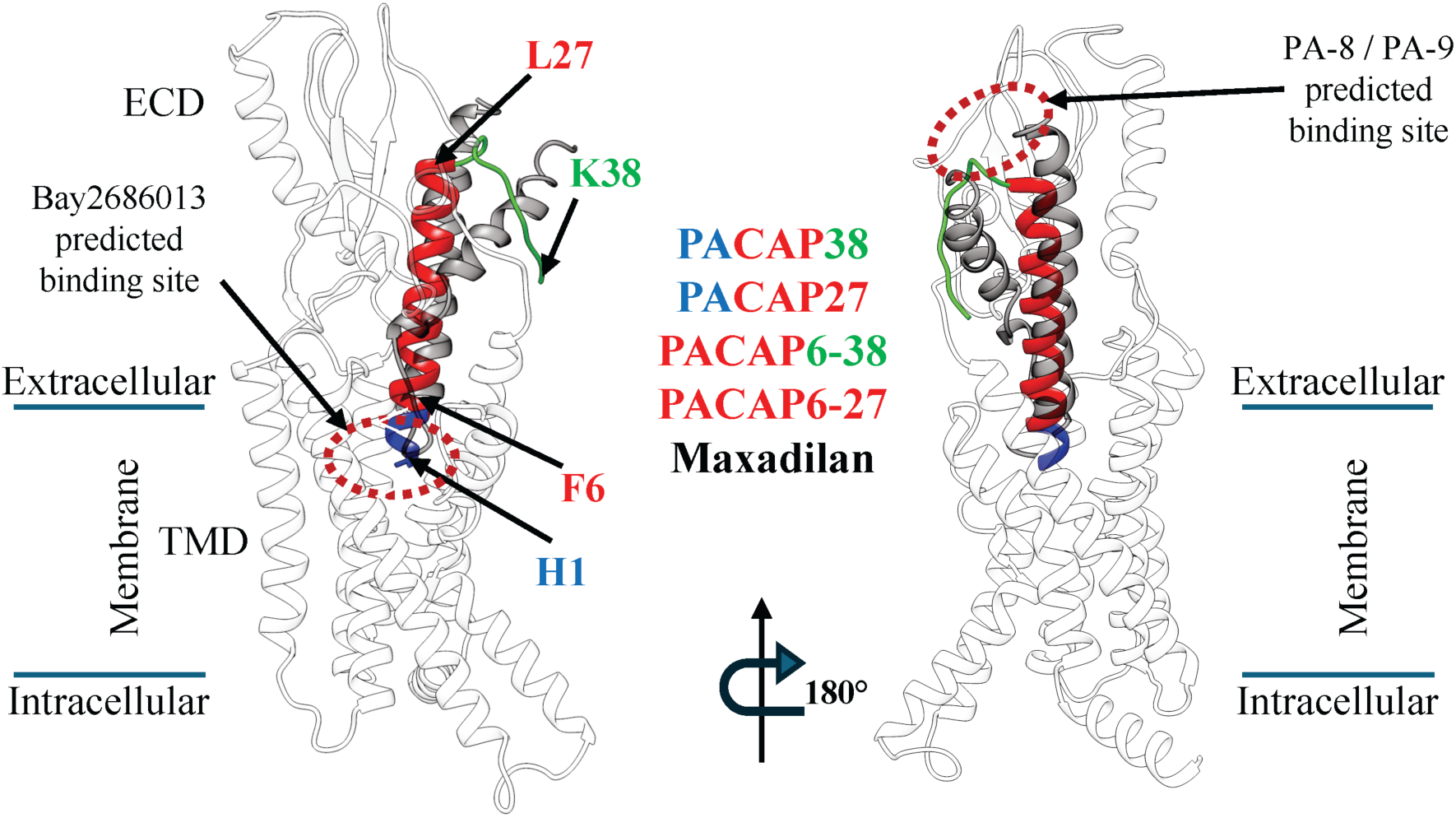
Binding regions of PAC1 ligands: agonists Maxadilan, PACAP38, and PACAP27 and their N-truncated forms, which act as antagonists (PACAP6-38 and PACAP6-27). Two side views of PAC1 (white ribbon), both from the cytoplasmic membrane (extra- and intracellular boundaries indicated). PACAP38 and PACAP27 N-terminal residues 1-5 (blue ribbon) are necessary for agonist activity; residues 6-27 (red ribbon) are common to both agonists and antagonists; only PACAP38 and PACAP6-38 have the C-terminal residues 28-38 (green ribbon). PAC1 ligand names are color-coded according to the ribbon coloring scheme. Maxadilan is shown in gray ribbon. Predicted binding sites for Bay268013 (TMD) or PA-8 and PA-9 (ECD) are indicated by red dashed ovals.

### 3.2. High level expression of hPAC1 in HEK293 cells has high potency PACAP signaling but limited sensitivity for peptide antagonist inhibition of responses

We developed a cellular PACAP inhibitor screen based on a clone of HEK293 cells expressing a split-luciferase-based cAMP sensor (HEK293_CBS), into which functional GPCRs of all classes could be expression-cloned using a retroviral vector with a selectable marker. Due to the ease of transduction of GPCR-expressing retroviral vectors into mammalian cell lines [30], it was possible to generate cell lines expressing human PAC1 (hPAC1), closely-related receptors such as VPAC1, VPAC2, and GLP1R, and unrelated GPCRs [30] to establish selectivity among potential assay hits. The HEK293_CBS_hPAC1 cell line was highly specific for activation by PACAP, compared to the related peptide VIP (Figure 2A**, right panel**), and was highly responsive to P38 and P27 with similar potency and efficacy. In comparison, the parental cell line HEK293_CBS showed minimal response to either PACAPs or VIP (less than 5% of the level for HEK293_CBS_hPAC1) and the potency of PACAP was similar to VIP (Figure 2A**, left panel**). Both HEK293_CBS and HEK293_CBS_hPAC1 cells displayed similar increases in cAMP accumulation in reponse to forskolin activation. The HEK293_CBS_hPAC1 line was used to perform an HTS assay of 156,059 compounds from the Molecular Libraries Probe Center Network compound library described previously [51]. In our assay, no inhibitory activity was observed for two SMOLs, an analog of PA-8 and the second an exact match to PA-9 (Figure 2B). This contrasts with the previously reported potent inhibition of P38-induced cAMP generation, and CREB phosphorylation, by these compounds in PAC1-expressing CHOK1 cells [20]. We therefore attempted further validation of our assay using the well characterized peptide inhibitors, P6-38 and M65. As shown in Figure 3A, neither peptide, in concentrations up to 10 µM, blocked P38- induced cAMP accumulation in HEK293_CBS_hPAC1 cells. P6-38 was also unable to antagonize P38-induced responses in these cells when cAMP concentration was measured directly by ELISA rather than using the cAMP biosensor read-out (Figure 3B). The highest concentrations of P6-38 or M65 tested were well above those employed for antagonism of PAC1 in neuroendocrine (NS- 1) cell-based biological assays (Figure 3C; [52]) or administered *in vivo* [53–60]. Due to the limited sensitivity of HEK293_CBS_hPAC1 cells to inhibition of P38-stimulated responses by well characterized peptide inhibitors, an alternate PAC1 assay system was pursued prior to further evaluation of putative hits from the HTS.

**Figure 2.**
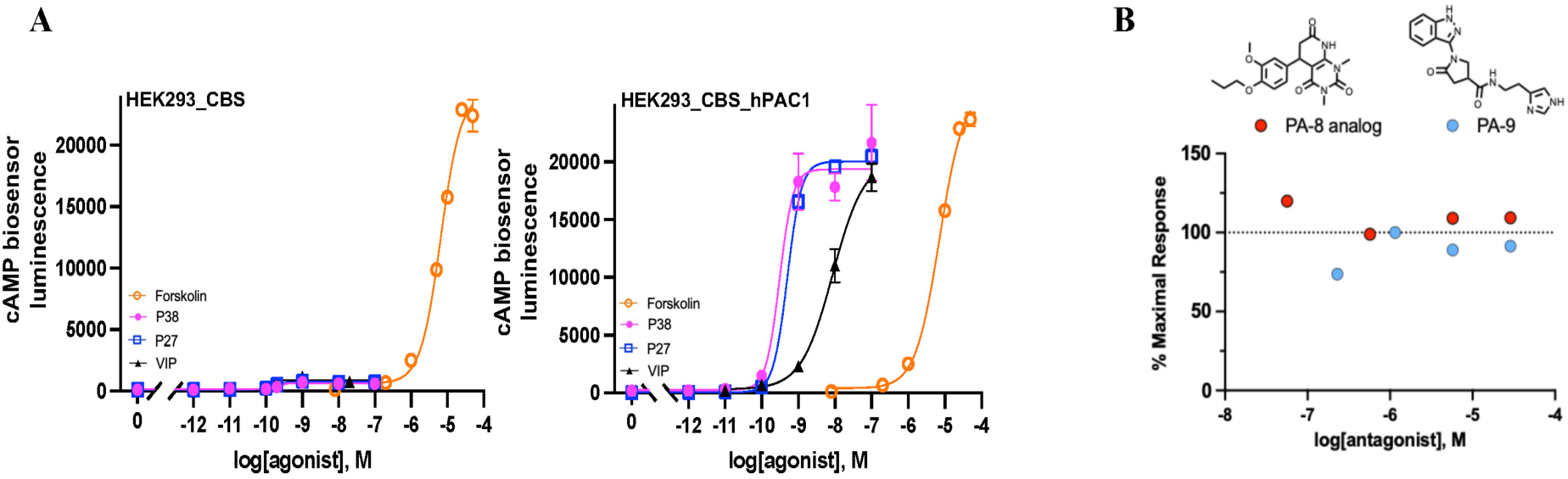
Establishment of a HEK293_CBS_hPAC1 cell line expressing exogenous cAMP biosensor (CBS) and hPAC1 to screen for antagonism of P38-induced cAMP elevation. **(A)** Characteristics of HEK293_CBS_hPAC1. Concentration response curves for PACAP27, PACAP38, VIP and forskolin on HEK293_CBS (left) and HEK293_CBS_hPAC1 (right). Cells were pretreated with isobutyl-1-methylxanthine (IBMX), an inhibitor of phosphodiesterase, to enhance cAMP detection. Luminescence signal was used as a readout (Y axis) to measure intracellular cAMP level in live cells after D-Luc pretreatment. Data are shown as mean ± SEM of triplicate repeats from a single experiment, which was repeated twice with similar results. **(B)** Effects of PA-8 analog and PA-9, contained in the NCATS HTS library, on P38-induced cAMP elevation in HTS screen. The % response reflects cAMP levels induced by antagonist plus P38, normalized to the maximal cAMP elevation elicited by P38 alone (set as 100%).

**Figure 3.**
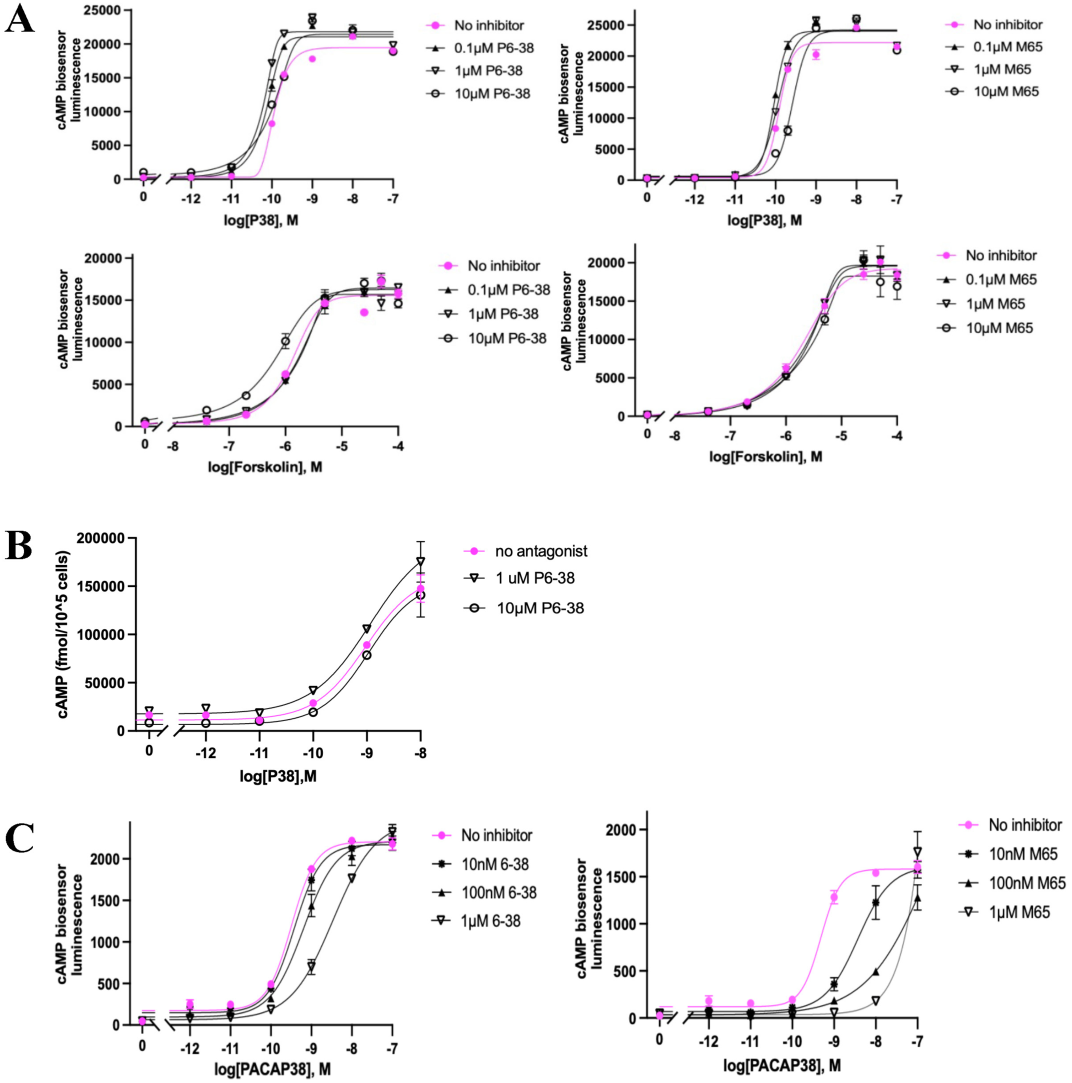
PACAP-induced cAMP in HEK293_CBS_hPAC1 cell line: effects of P6-38 and M65. **(A)** P6-38 or M65 addition, in P38-induced cAMP assay in HEK293_CBS_hPAC1 cells with measurement of cAMP-induced bioluminescence. Cells were pretreated with PAC1 antagonist P6-38 (left panel) or M65 (right panel) for 10 minutes at the concentrations of 0.1 µM, 1 µM, and 10 µM prior to addition of P38 (top panel) or forskolin (bottom panel) to activate cAMP production. **(B)** Direct cAMP ELISA assay confirms that P6-38 does not inhibit P38-induced cAMP elevation in HEK293_CBS cells exogenously expressing hPAC1. ELISA assay was used to directly measure cAMP concentration in a 96-well plate with cAMP antibody (Arbor Assays). cAMP levels measured after cell lysis are expressed as fmol per10⁵ cells on the ordinate. (C) An endogenously expressed PAC1 in rat neuroendocrine NS-1 cells can be activated by P38 and its activation is inhibited by P6-38 or M65 using the CBS assay after expressing CBS in NS-1 cells. The concentration response curve of P38 or forskolin without any antagonist is shown in pink. The data shown above represent the averages from three replicates (n = 3), with standard errors of the mean indicated by vertical bars. The experiments were independently repeated at least three times with qualitatively and quantitatively similar results.

### 3.3. Neuroblastoma SH-SY5Y cells endogenously xpressing hPAC1 and engineered to express CBS have an improved sensitivity for detection of PAC1 antagonists

Lack of inhibition of PACAP action by known peptide antagonists in the PAC1-expressing HEK293 cells motivated a search for a neuroendocrine cell line with appropriate sensitivity to PAC1 inhibition by peptide antagonists and appropriate properties for a high-throughput antagonist assay. NS-1 cells do not fulfill the criteria required for high-throughput screening, due to poor adherence in the absence of substrate. Therefore, we examined SH-SY5Y human neuroblastoma cells, a cell line that has been noted to express functional PAC1, mainly PAC1null [61], which is the isoform most highly expressed in human and rodent brain [62], and respond to P27 and P38 [61]. We installed the split-luciferase cyclic AMP biosensor in SH-SY5Y cells (SY5Y_CBS) and established that SY5Y_CBS cells respond to P38 in a concentration-dependent manner. This response was also concentration-dependently inhibited by P6-38 and M65, which by themselves had no effect on forsokolin-induced cAMP elevation in this assay (Figure 4A). The inhibition of the response to P38 in SY5Y_CBS cells was further confirmed using a direct cAMP ELISA assay (Figure 4B). P27 and P38 were equipotent in this cell line (EC_80_ ∼0.2 nM), with VIP about 100 times less potent (EC_80_ ∼20 nM), as expected for PAC1 activation in these cells (Figure 5A). We next pursued an HTS protocol to identify PAC1 antagonists, employing SY5Y_CBS cells. P6-38 was more potent against P27 (IC_50_ ∼20 nM) than against P38 (IC_50_ ∼100 nM), as estimated from Figure 5B. We therefore chose P27 as the agonist in the HTS assay. Using P27 to drive PAC1-dependent cAMP accumulation, we screened 3,008 compounds, including 192 compounds picked from previously investigated HTS HEK293_CBS_hPAC1 assay and 2,816 compounds from the HEAL library [32]. Here we also counterscreened compounds with inhibitory activity against P27 using forskolin-stimulated responses as a control for non-receptor mediated inhibition. 25 compounds were selected that inhibited P27-induced cAMP accumulation, at concentrations between 5 µM and 30 µM, but exhibited less potency to inhibit forskolin-induced cAMP generation. Five of these 25 compounds were also able to inhibit P38-induced cAMP elevation when tested under 96-well format assay conditions. The chemical structures of these compounds are shown in Figure 5C. However, we were unable to confirm selective activity for P38 compared to forskolin inhibition in the 96-well SY5Y_CBS assay (Figure 5D), an important corrollary for specificity and locus of action of putative PAC1 antagonist compounds.

**Figure 4.**
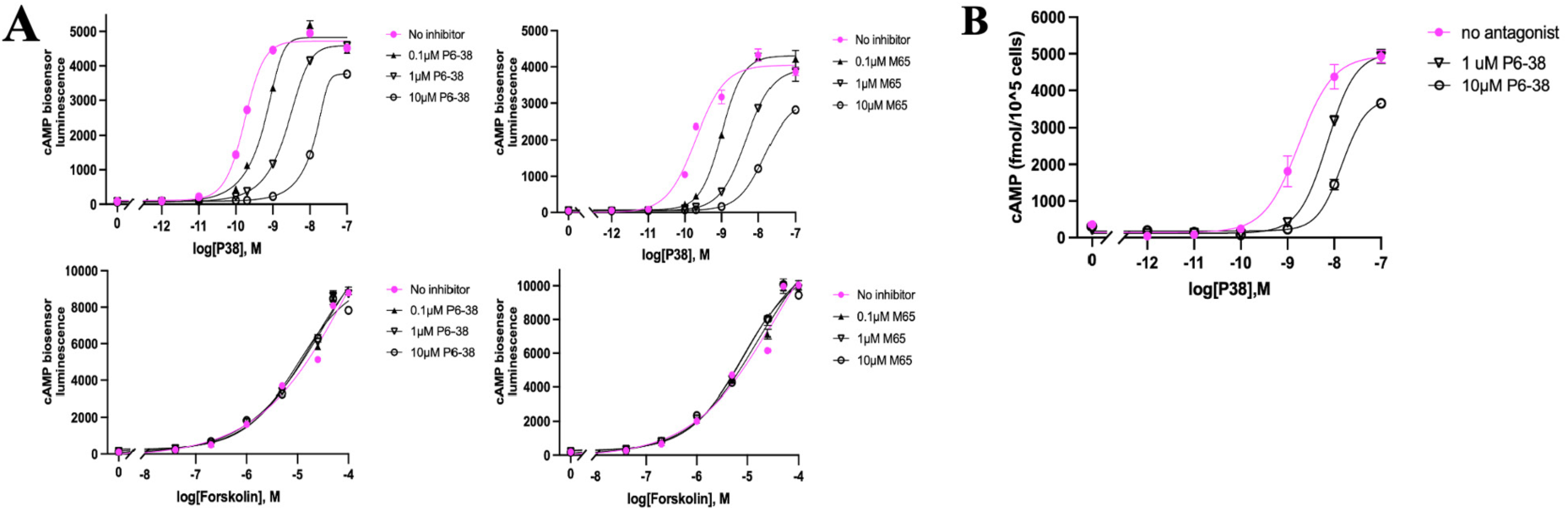
Elevation of cAMP by P38 in SY5Y_CBS cells: dose response curve and inhibition by P6-38 and M65. **(A)** The antagonist effect P6-38 (left panel) and M65 (right panel) on the cellular cAMP response after P38 (top panel) or forskolin (bottom panel) induction was detected by cAMP CBS assay. **(B)** Direct cAMP ELISA assay confirms that P38-induced cAMP elevation is inhibited by P6-38 in SY5Y_CBS cells. Data are presented as mean ± SEM from a representative experiment performed in triplicate. Error bars are shown; where not visible, they are contained within the symbols. Similar results were obtained in three separate experiments.

**Figure 5.**
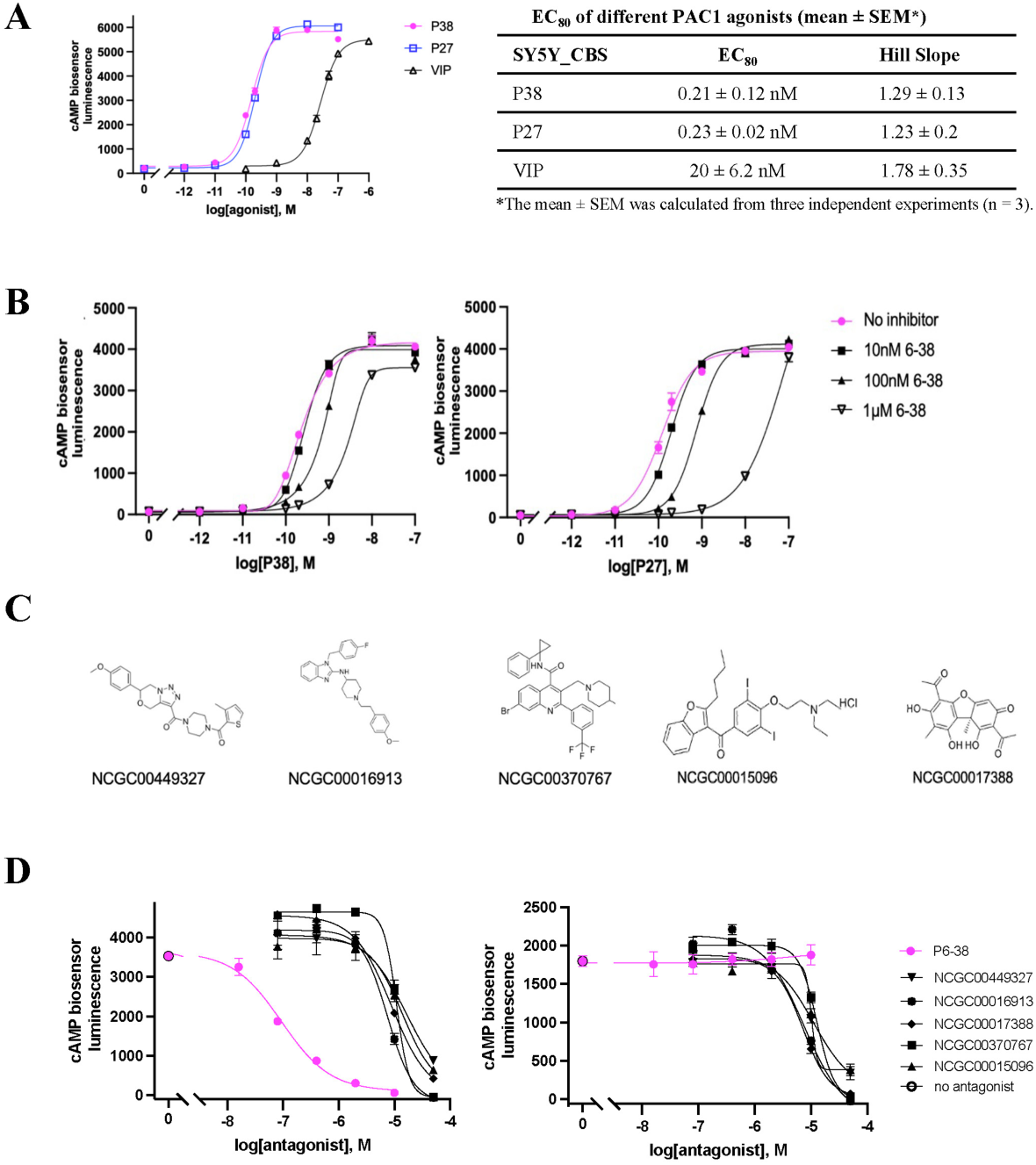
Screening of 3,008 compounds as PAC1 antagonist candidates. Using P27 or P38 at a concentration of 0.2 nM (EC_80_) to screen. **(A)** P38 and P27 are equipotent agonists in SY5Y_CBS cells when measured in the CBS assay. VIP is a less potent agonist. **(B)** P6-38 can block both P27- and P38-induced cAMP elevation in SY5Y_CBS cells but is a more potent antagonist against P27 when tested in the CBS assay. **(C)** Structures of the five compounds that showed activity against P38 in SY5Y_CBS 96- well counterscreen, from the 25 hits identified after initial screening of 3,008 compounds from HEAL library and NCATS collection for PAC1 antagonism using SY5Y_CBS cells in HTS format with P27 as the agonist. **(D)** Testing the five compounds in C for antagonist effect on P38- and forskolin-induced cAMP elevation using the CBS assay. Data are presented as the average value of three replicates from a representative experiment. Error bars represent the standard error of the mean (SEM); where not visible, they are contained within the symbols. Three independent experiments were performed with similar results.

### 3.4. Evaluation of published small molecule PAC1 antagonists in SY5Y_CBS cells

We next tested previously reported PAC1 antagonists, including PA-8 and related compounds [20, 21, 25, 26, 63, 64] for their PAC1 antagonist effect and specificity using SY5Y_CBS cells, in comparison to P6-38 as the positive antagonist control. Compounds Beebe-1 and Beebe-2 (Figure 6A) correspond to the two lead hydrazides 1 and 2 identified by Beebe et al. as PAC1 antagonists based on inhibition of labeled P27 binding in an *in vitro* assay system [25]. Both SMOLs inhibited cAMP accumulation by P38 less than 50% at 10 µM. However, inhibition was not specific for PAC1, as cAMP production by forskolin was inhibited to the same extent at the corresponding concentration (Figure 6A). When PA-8 and PA-9, and derivatives 2o and 3d/PA-915, were tested in SY5Y_CBS cells, inhibition of PACAP action was not detectable, relative to P6-38, either using the CBS assay, or direct cAMP ELISA assay (Figure 6B-D). We observed an inhibiton of PAC1 activation by P38 by over 50% at the concentration of 10 µM with the recently reported putative PAC1 antagonist compound Bay2686013 [26]. The potency of Bay2686013 to inhibit P38 activation of PAC1 was lower than that of P6-38 by approximately two orders of magnitude (Figure 6C-D).

**Figure 6.**
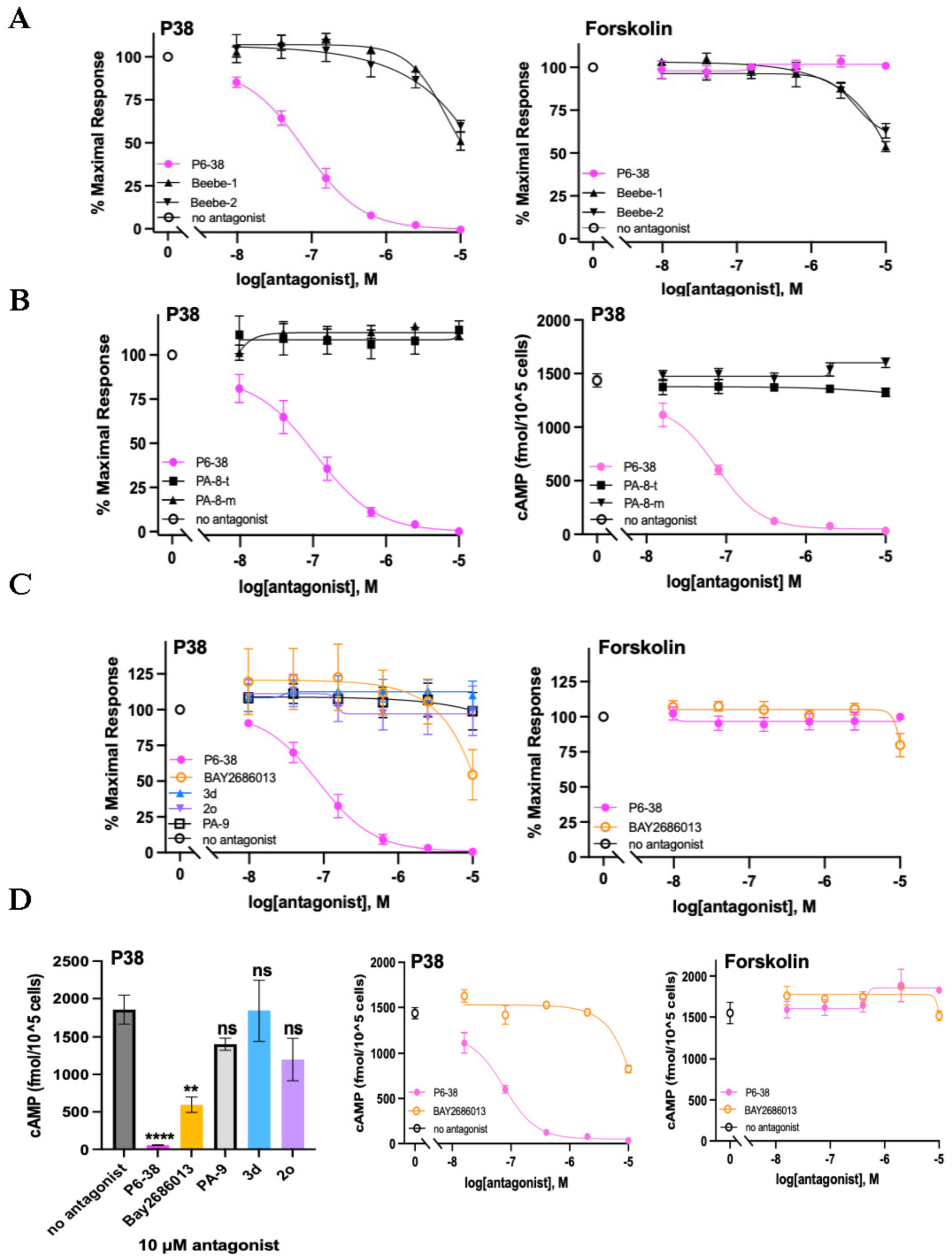
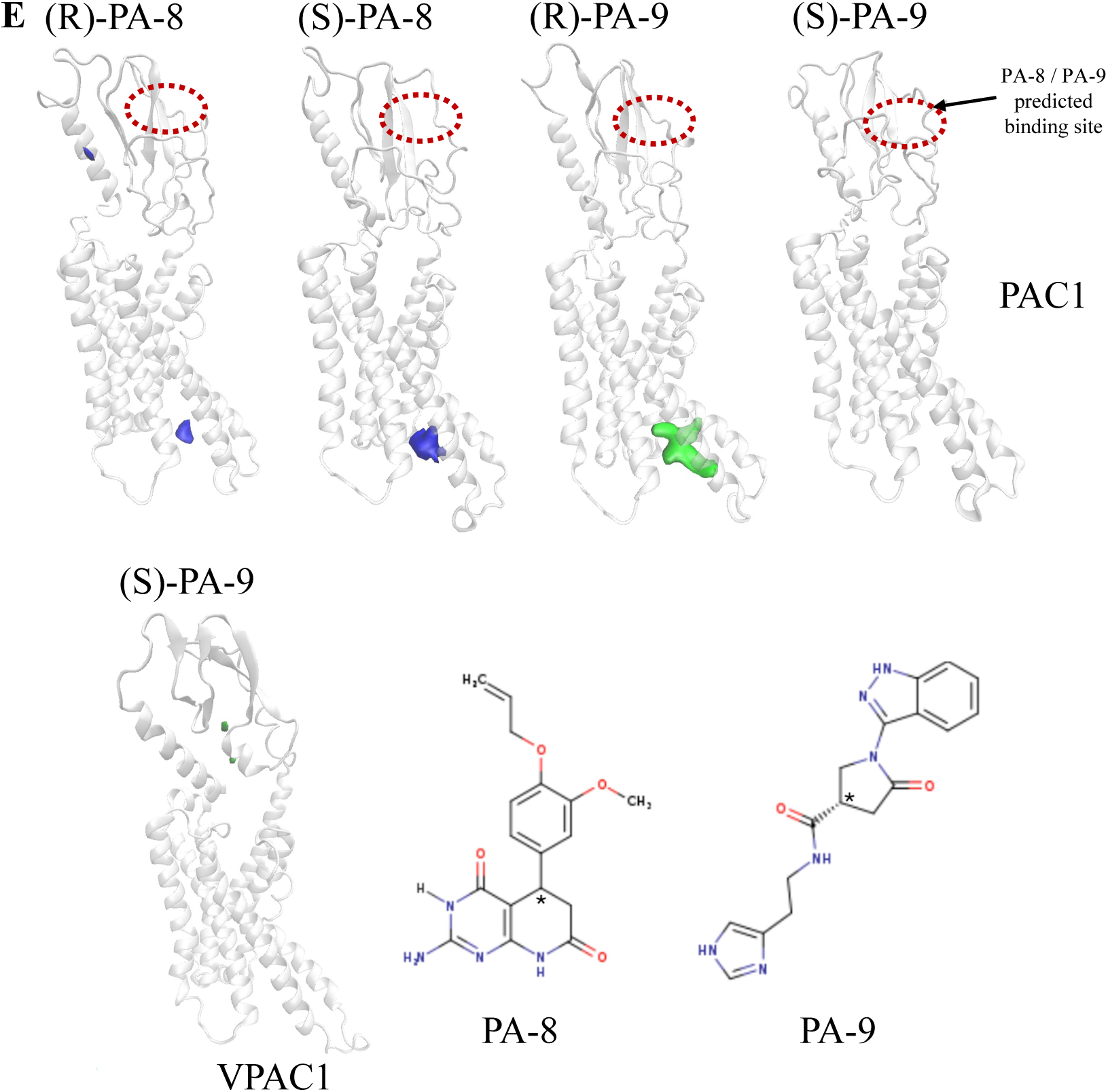
Evaluation of previously reported small molecule PAC1 antagonists in SY5Y_CBS cells. **(A)** Effects of Beebe-1 and Beebe-2 on P38- and forskolin-induced cAMP elevation when tested in the CBS assay. **(B)** Effects of PA-8 obtained from two different sources (t:Tocris, m:MedChemExpress) on P38-induced cAMP accumulation when tested in the CBS assay (left panel) or direct cAMP ELISA assay (right panel). **(C)** PA-8 derivatives: PA-9, 3d, 2o, and Bay2686013 were tested for inhibition of P38-induced cAMP elevation in the CBS assay (left panel), and compounds inhibiting P38 tested for inhibition of forskolin (5 µM) action (right panel). (D) Direct cAMP ELISA assay to confirm results of CBS assay in 5C. Left panel, Middle panel, all treated with 1 nM P38. Left panel: The values treated with the antagonist (10 µM) were compared to the values without any antagonist for significant differences using one-way ANOVA followed by Dunnett’s post-hoc test. Data are expressed as mean ± SEM. Significance levels are indicated as follows: ns (not significant), **P<0.01, ****P<0.0001). Middle panel, dose-responsive inhibition of P38 response. Right panel, inhibition of forskolin response (25 µM) . Data displayed in CBS assays represent average values from three individual experiments. ELISA assays shown were repeated with similar results. **(E)** Volumetric maps of the multiligand MD simulations do not suggest binding of the small molecules PA-8 and PA-9. Different enantiomers of PA-8 and PA-9 were simulated in separate systems. VPAC1R was simulated with (S)-PA-9 as a negative control (not inhibited by PA-9 in [20]. Volumes characterized by an occupancy of SMOL atoms ≥ 25% (of the total simulation time) are shown in blue (PA-8) and green (PA-9). No persistent interactions were sampled either at the predicted [20] ECD binding sites (dotted ovals) or other ECD positions. The region of PAC1 most prone to interact with PA-8 or PA-9 was the intracellular G_s_ protein binding site, as indicated by colored volumes for (R)/(S)-PA-8 and (R)-PA-9. This result is likely an artifact due to the active PAC1 state used for the simulations, which presents the G_s_ protein binding site in open conformation.

Microsecond molecular dynamics (MD) simulations of PA-8 and PA-9 binding to PAC1n did not produce persistent PAC1:SMOLs interactions in the predicted ECD binding site for these ligands (Figure 6E). The total MD sampling time (56 µs for PA-8 and 51 µs for PA-9, as total simulation time multiplied by the number of SMOL molecules in the simulation box) should have allowed for at least one SMOL to bind the solvent-accessible binding sites on the ECD [11]. Similar results were observed using the same simulation approach on VPAC1, which was reported as a non-binder of PA-8 or PA-9 [20]. PA-8 and PA-9 transiently engaged PAC1 at the intracellular G_s_ protein recognition site. However, such interactions are likely an artifact due to the use of an active state receptor for the simulations but with the G protein removed, which has a wide intracellular cavity that should not be present in the apo, inactive PAC1 responsible for PA-8 and PA-9 binding. A caveat on the current empirical and simulated observations that lack evidence of productive engagement of the SMOLs at PAC1, is that these studies predominantly used the most common isoform, PAC1n. It is possible that these ligands have higher activity at other isoforms.

In observing limited docking in our simulations for SMOLs reported to block PACAP activation of PAC1 (Figure 6E), we hypothesized that re-examination of peptide-based inhibition, based on current structural information [13, 19] and the ability to examine PAC1 engagement in a battery of high-content cell-based assays, such as the SY5Y_CBS assay employed here, was justified.

### 3.5. Peptide modifications that enhanced antagonist activity of secretin peptides at their cognate receptor did not improve PACAP inhibition of PAC1

Lactam modifications can enhance the efficacy of secretin receptor antagonists by stabilizing specific conformations, such as α-helical structures, which are crucial for receptor binding. In a study by Dong and colleagues [65], incorporation of a lactam bridge between residues 16 and 20 in secretin analogues resulted in a 22-fold increase in binding affinity compared to the unconstrained peptide. Further, substitution of Leu residue 22 with the bulkier and more sterically constrained amino acid, L-cyclohexylalanine (Cha), further enhanced the binding affinity and improved the antagonist effect of secretin 5-27. Given the structural similarities between the secretin and PACAP receptors, it was plausible that lactam modifications could similarly enhance PACAP binding. This hypothesis was supported by MD simulations with P6-27 peptide analogs with a lactam derivatization (P6-27Lb) and a Cha substitution at Leu23 (P6-27Lbx1) or Leu27 (P6-27Lbx2) that decreased peptide dynamics, indicative of tighter interactions (root mean square deviations, RMSD, values in Figure 7B). We therefore substituted the corresponding residues in P6-27 to form a lactam bridge (Figure 7A) and tested its antagonist effect on PAC1. Contrary to expectation, the lactam bridge and Cha substitutions decreased the antagonist potency of P6-27 with P27 as an agonist (Figure 7C).

**Figure 7.**
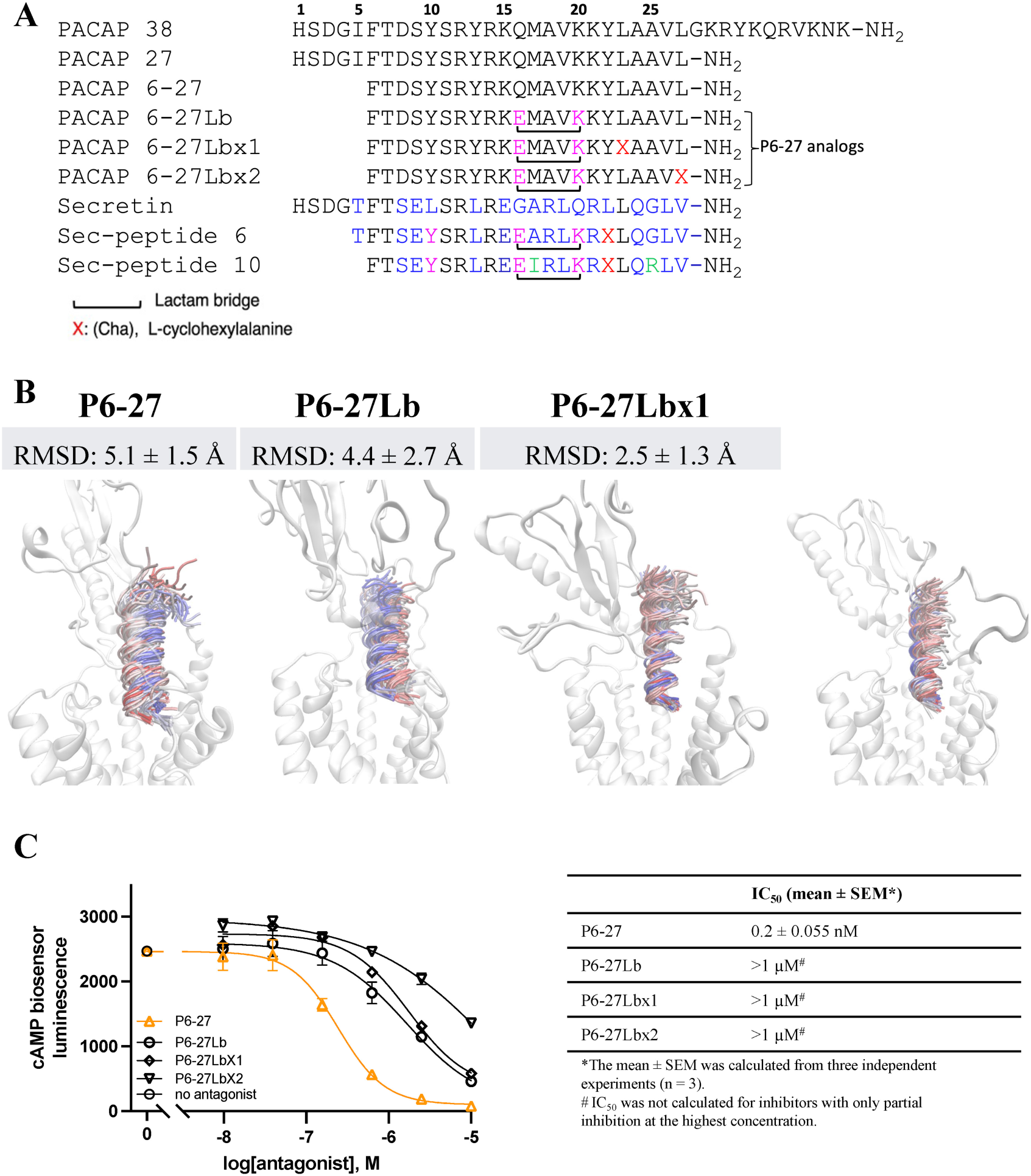
Modifications to P6-27 increase simulated engagement stability but decrease antagonist potency at PAC1. **(A)** Amino acid alignment of PACAP with secretin and the substitutions in P6-27 to form a lactam bridge with or without Cha modification at the position corresponding to secretin antagonist analogues (Sec-peptide 6: (Y^10^,c[E^16^,K^20^], Cha^22^)sec(5–27) and Sec-peptide 10: (Y^10^,c[E^16^,K^20^],I^17^,Cha^22^,R^25^)sec(6–27). The amino acids in pink font are conserved among P6-27 analogs and secretin analogues but differ from natural secretin sequences. Amino acids in blue font are conserved in natural secretin sequences. Amino acids in black font are conserved in PACAP sequences. X: L-cyclohexylalanaine; black brackets: Lactam bridge. **(B)** Backbone stability (RMSD) of P6-27 and derivatives during MD simulations of binding to PAC1. The peptide backbone is colored according to the simulation time of three merged MD replicas, from red (first MD frame) to blue (last MD frame), plotted every 1 ns. **(C)** P6- 27Lb, P6-27Lbx1, and P6-27Lbx2 did not increase the inhibitory effect of P6-27 against 0.2 nM P27 (EC_80_) when tested in the CBS assay. Graph shown is one of the three individual experiments to calculate the IC_50_ in the table. The error bars in the graph are SEM calculated from three replicates in that experiment, when the error bars are not shown, these are contained within the symbols.

A comparison of P6-27 in the SY5Y_CBS assay revealed a pronounced differential potency for blockade of P27 compared to P38 (Figure 8). The identical potency of P27 and P38 for PAC1 binding and activation would imply that the contact points for both peptides were common to, and restricted to, the first 27 residues. However, the lack of equipotency of P6-27 against P27 and P38 implies a role for residues beyond 1-27, in P38, for high-affinity binding to PAC1, as suggested by the initial structure-activity profiling of P6-38 [14].

**Figure 8.**
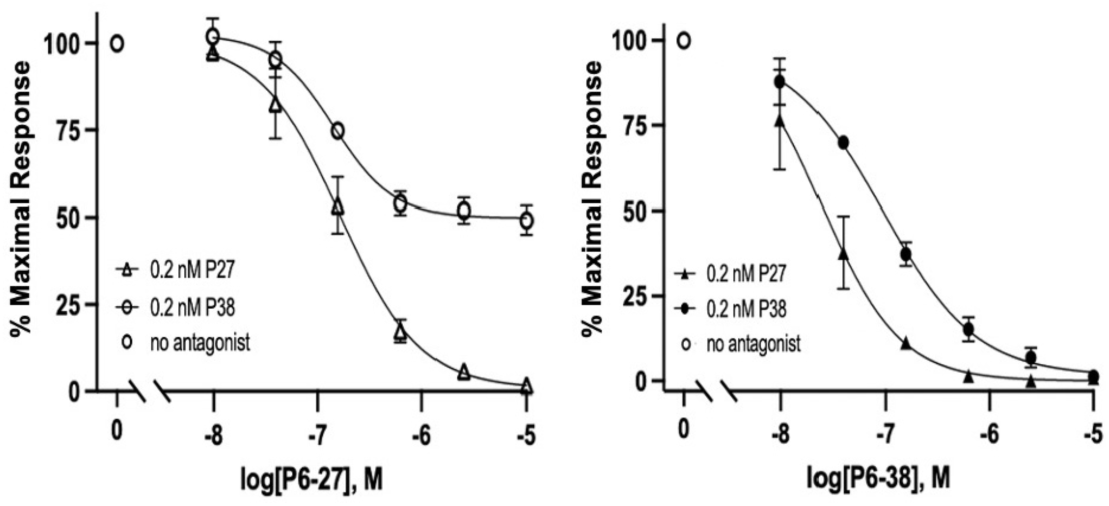
P6-27 inhibition of PACAP-stimulated cAMP elevation. P6-27 inhibition of PACAP-stimulated cAMP in SY5Y_CBS cells. P6-27 blocks P27-stimulated (0.2 nM) cAMP elevation (triangles) with an IC_50_ of ∼0.1 μM, and blocks P38-induced (0.2 nM) cAMP elevation (circles) to only 50% at a concentration of 1 μM and remains at ∼50% until the highest concentration tested (10 μM) in the CBS assay. Data are shown as mean ± SEM of 3 independent experiments. Where error bars are not visible, they are contained within the symbols.

The differential effect of the inhibitor peptides on P27- versus P38-mediated responses is consistent with distinct modes of binding for the two agonist peptides, despite being equipotent in the cAMP accumulation assays. This implies that additional important receptor interactions occur through residues of P38 beyond residue 27, leading us to undertake MD simulations of the binding of P6-38 without modification and peptide analogs with a lactam derivatization (P6-38Lb), or lactam derivatization combined with a Cha residue in place of Leu23 (P6-38Lbx1) (Figure 9A and B). As noted above, such modifications could improve receptor-compatible conformations, as predicted and found with the highly related secretin and its receptor [65, 66]. Molecular modeling suggested stability similar, or improved, for P6-38Lb and P6-38Lbx1 residues 6-31 compared to P6-38 (Figure 9A). As shown in Figure 7C **and 9C**, introduction of a lactam bridge alone reduced the inhibitory potency of both P6-27 and PA6-38, although the combination with Cha in P6- 38Lbx1 maintained equivalent potency for the longer peptide (Figure 9C), distinct from the impact on the P6-27 (Figure 7C). Nonetheless, the failure to improve inhibitory potency of the antagonists indicated that secretin- secretin receptor (SCTR) interaction is not a pertinent model for design of PACAP-based inhibitors leading us to focus on understanding the interactions driving binding stability through detailed analysis of our MD data with P6-38.

**Figure 9.**
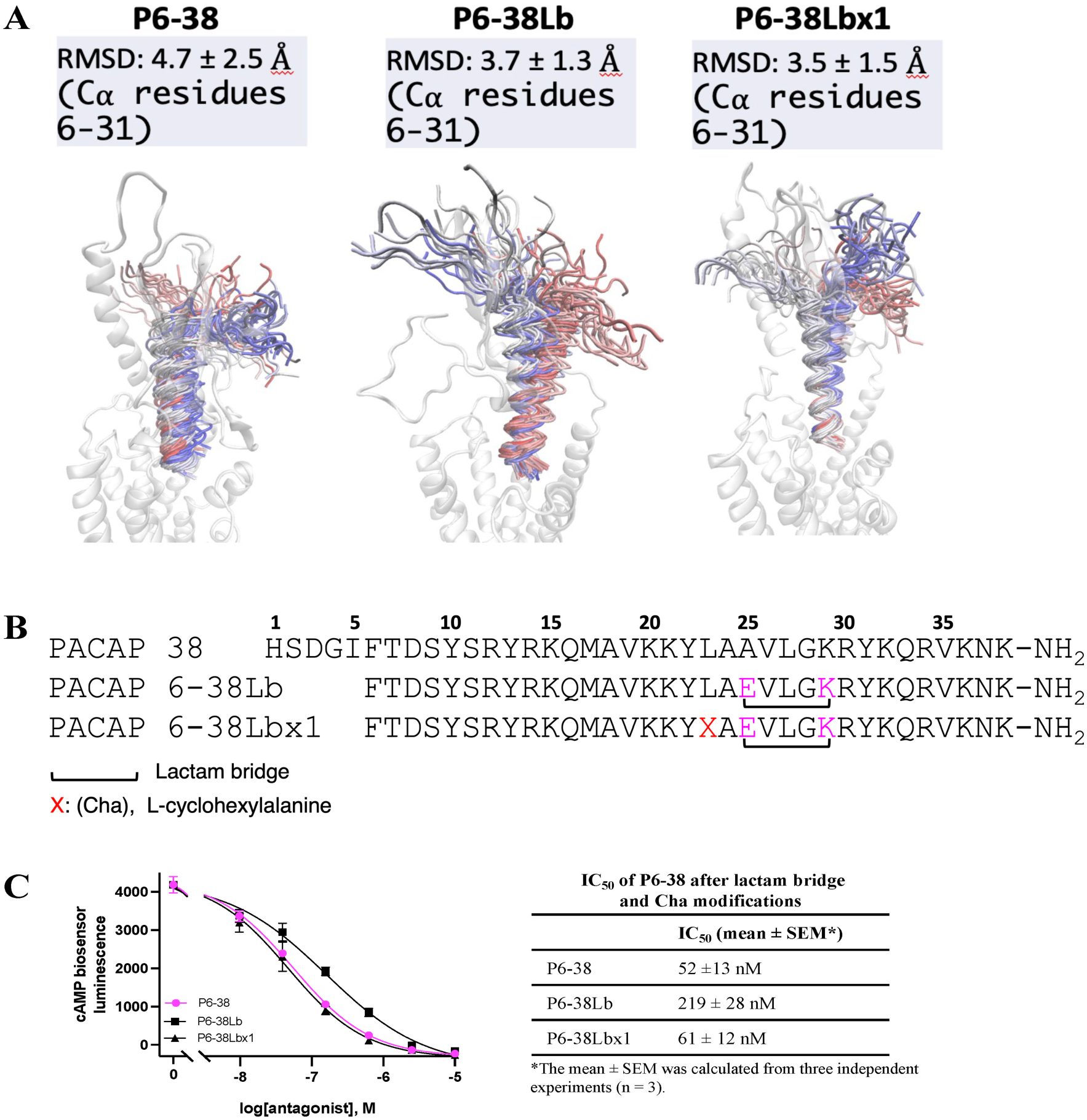
Testing the effects of lactam bridge and Cha modification of P6-38 on its antagonist effect. **(A)** Molecular modeling results of modifying P6-38 to create a lactam bridge interaction to extend the C-terminal helix of P6-38 to improve its binding interactions with the ECD of PAC1. The peptide backbone is colored according to the simulation time of three merged MD replicas, from red (first MD frame) to blue (last MD frame), plotted every 1 ns. **(B)** Amino acid sequences of the proposed peptides are shown (X: L-cyclohexylalanine; black bracket: Lactam bridge). **(C)** Antagonist potency of P6-38Lb and P6-38Lbx1 was tested in the CBS assay using 0.2 nM P38 as the agonist . The graph represents one of three independent experiments. Error bars indicate the SEM of three technical replicates from the representative experiment.

### 3.6. Molecular dynamics and cryo-EM data are consistent with P38 binding to PAC1 mainly via its first 30 residues, with residues 31-38 potentially dispensable for binding, which was confirmed using SY5Y_CBS cellular assay and NS-1 neuritogenesis assay

Detailed analysis of MD simulations of the binding of P6-38 to PAC1 indicated strong receptor contact between residues 28-30 in P6-38 but not beyond residue 30 (Figure 10A-E). R30, in particular, was predicted as pivotal for antagonist binding and stabilization at the ECD (Figure 10B-C). MD simulations of the C-terminal truncated P6-30 suggested persistent salt bridges and hydrogen bonds between R30 and two clusters of acidic residues in the receptor (Figure 10D-E), corroborating the importance of a positively charged residue in position 30. Hence, we designed, and tested in the SY5Y_CBS assay, a series of P6-38 C-terminal deletants (Figure 11A-B). Systematic deletion of residues revealed equipotency of P6-38 through P6-30. P6-29 through P6- 27 were able to act as antagonists only at concentrations of 1 µM or above, and achieved only partial inhibition at those concentrations (**Figure 11B**). These results were confirmed in direct cAMP assay by ELISA (**Figure 11C**).

**Figure 10.**
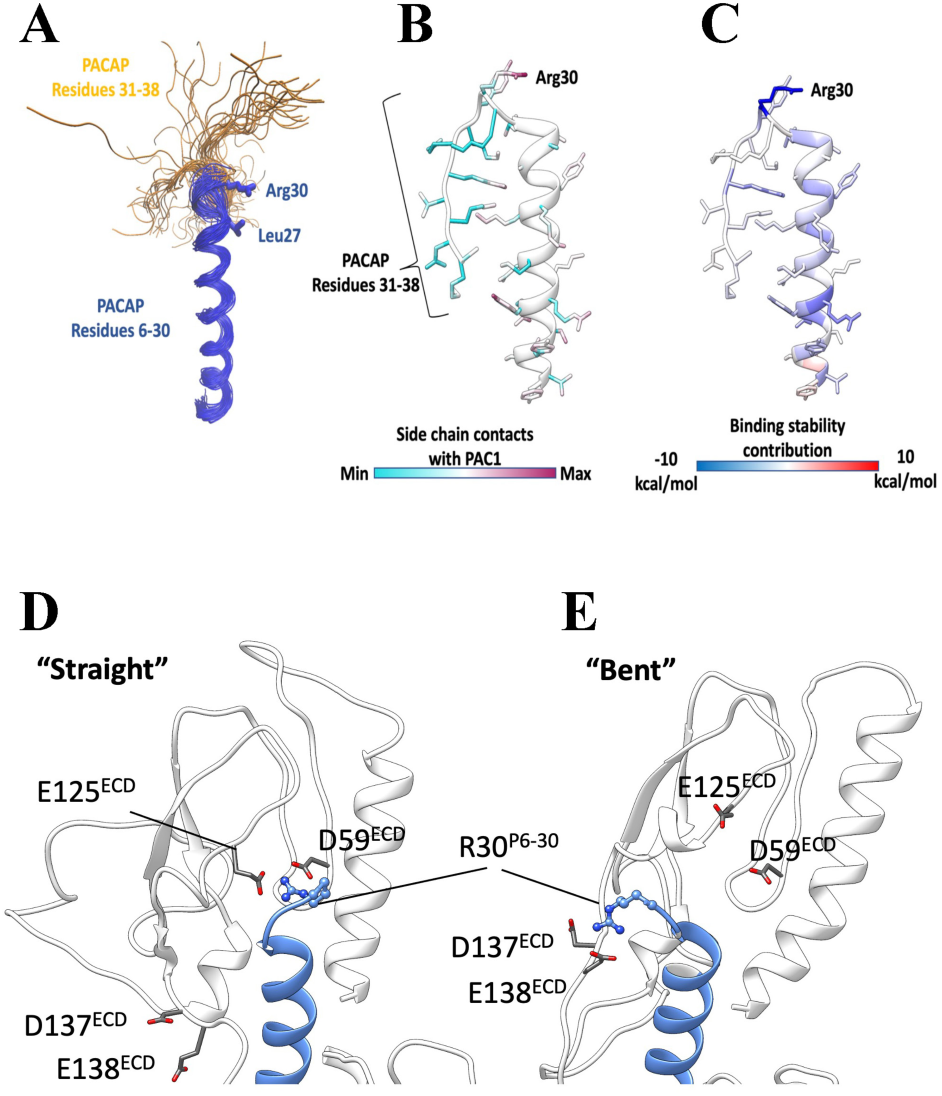
Molecular modeling of PAC1 antagonists P6-38 and P6-30. Molecular dynamics is consistent with P6-38 binding to PAC1 mainly via its first 30 residues, with residues 31-38 potentially dispensable for binding. **(A)** P6-38 superimposed snapshots from molecular dynamics simulations. The C-terminal residues 31-38 (shown in yellow) were highly flexible. **(B)** P6-38 residues 31-38 did not stably interact with PAC1. **(C)** P6-38 per residue contribution to computed binding stability: R30 is predicted to be the most stabilizing residue of the agonist C-terminal. **(D-E)** P6-30 interactions with acidic PAC1 residues in two different conformations. **(D)** In the “straight” conformation, P6-30 R30 hydrogen bonds with D59 and E125 of PAC1. **(E)** In the “bent” conformation, P6-30 R30 hydrogen bonds with D137 and E138 of PAC1. (PAC1, white ribbon with grey side chains; P6-30, blue ribbon with blue side chains).

**Figure 11.**
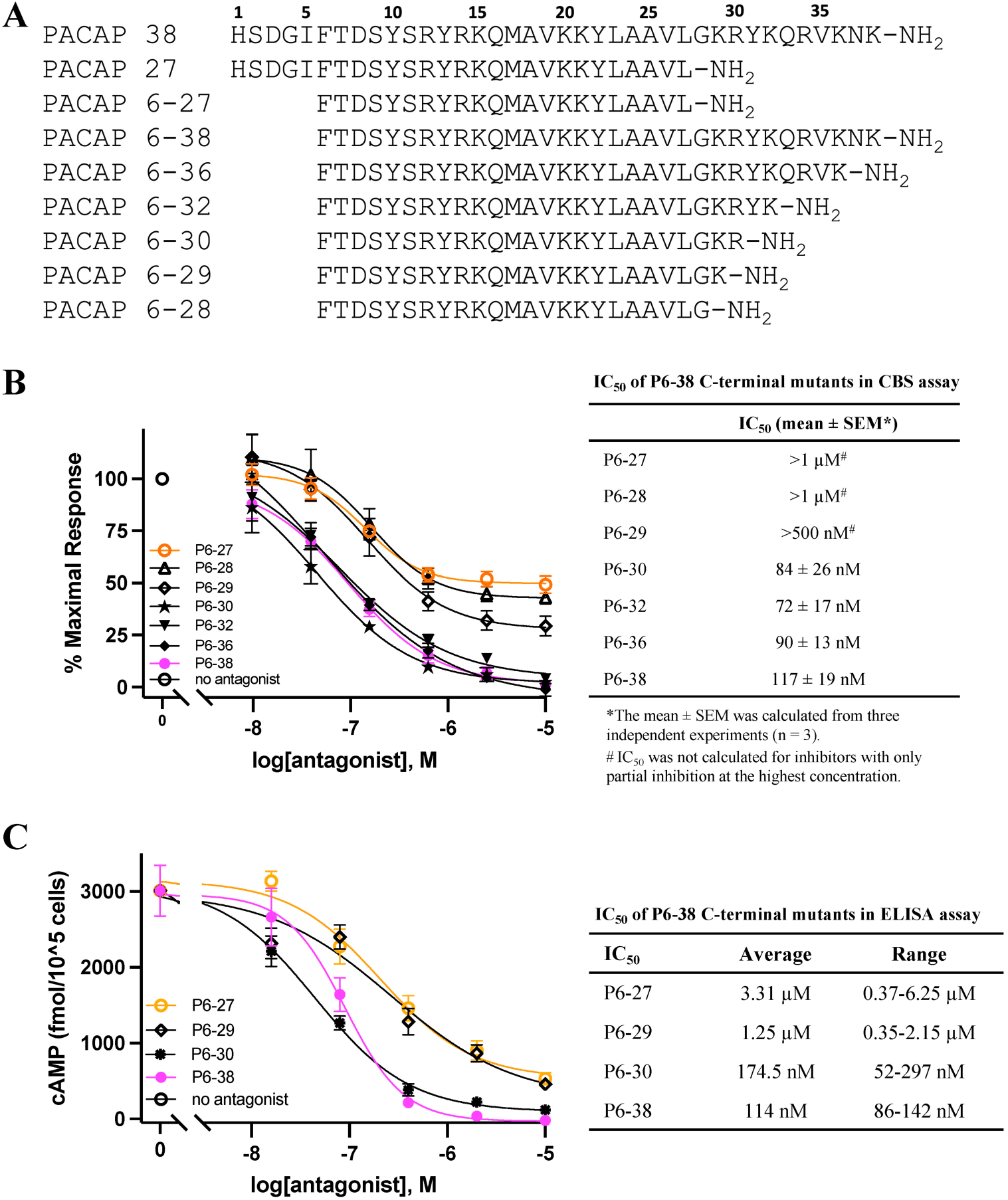
Investigation of P6-38 deletants as PAC1 antagonists. **(A)** Amino acid sequences of P38 and P6-38 and its deletants used in the current studies. Residue numbers correspond to numbering of P38. All peptides were amidated at the carboxy-terminal. **(B)** Effects of sequential deletion of C-terminal amino acid residues of P6-38 on inhibition of P38-induced cAMP elevation in SY5Y_CBS cells using CBS assay. 0.2 nM P38 (EC_80_) was used for cAMP stimulation. Three independent experiments were carried out to obtain the IC_50_ for each antagonist shown in the graph and table. **(C)** Direct cAMP ELISA assay confirms the differential effect of C-terminal terminal deletion at residue 30 on the antagonist effect of P6-38 to inhibit P38-induced cAMP elevation. 1 nM P38 was used in the ELISA assay. This experiment was repeated and the average and range of IC_50_ values from the two independent ELISA experiments are shown in the table. Graphs shown are from one representative experiment with mean ± SEM from three repeats in the corresponding experiment.

We then tested the ability of deletants of P6-38 to inhibit P38-induced neuritogenesis, a long-term effect of continuous exposure to PACAP over a period of hours to days [67]. The relative inhibition of P38-induced neuritogenesis induced by 10 µM P6-38, P6-30, P6-29 or P6-27 was similar to that determined in SY5Y_CBS cells (Figure 12), predicting likely translation of our findings to *in vivo* studies.

**Figure 12.**
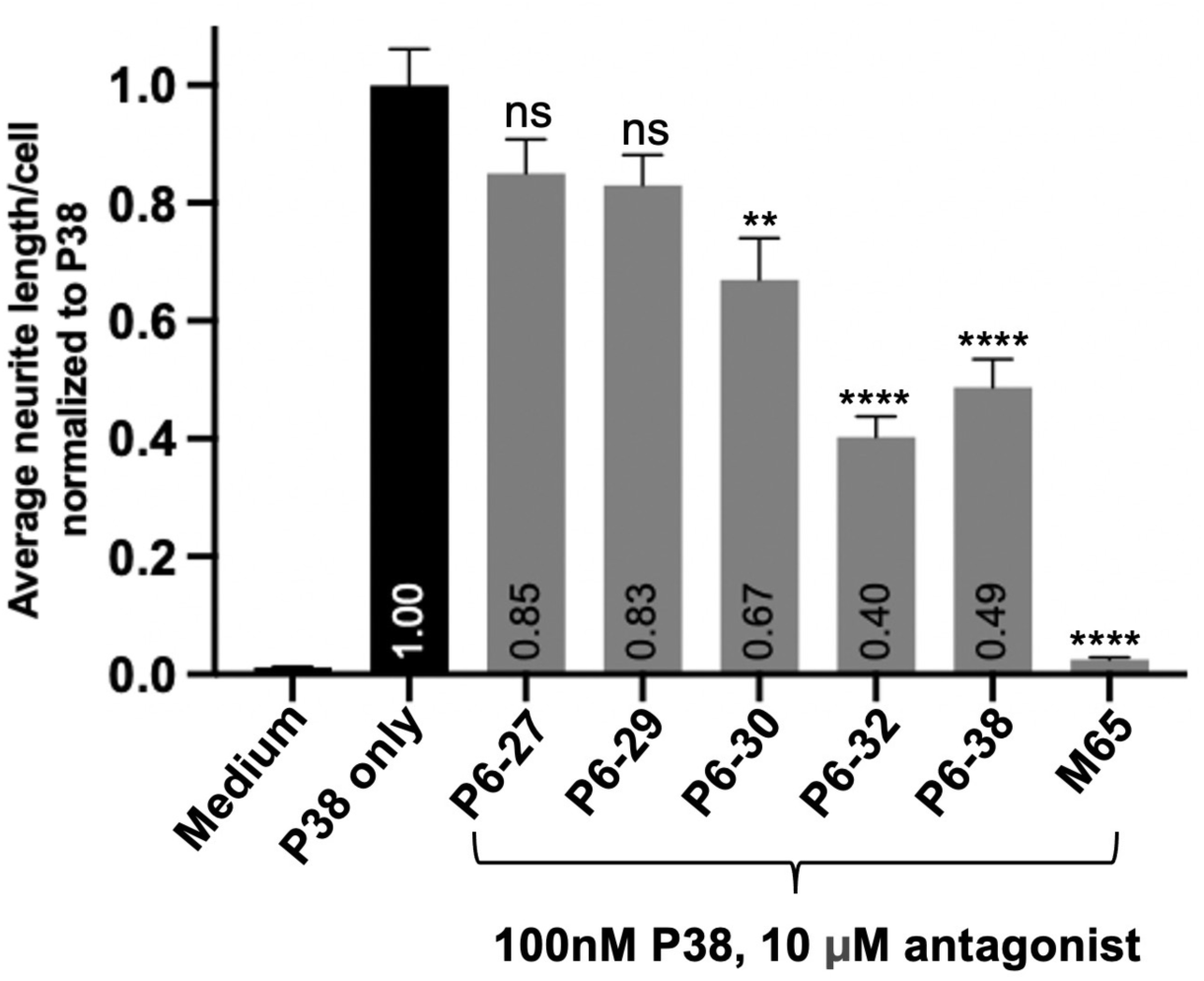
Inhibition of P38-induced neuritogenesis by P6-38 deletants in NS-1 cells. NS-1 cells were pre-treated with P6-38 deletants (10 μM) for 30 minutes before the addition of 100 nM P38. Images were taken at 48 hours after PACAP treatment. Neurite length and cell number were quantified using NIS-Elements AR Analysis with the AI Image Processing module. The bar graph shows the average (located within the bottom of the bar) from three separate experiments (n = 3), each performed in triplicate. The antagonist-treated values were compared for significant differences to P38 alone using one-way ANOVA with Dunnett’s post-hoc test, mean ± SEM, ns: not significant, *P>0.05, ** P<0.01, **** P<0.0001).

In summary, we describe here specific predictions for molecular features of P27, P38, and the antagonist peptide, P6-38, binding to PAC1, based on MD simulations built from cryo-EM structural data that are consistent with an affinity-trap model for binding and its antagonism by peptide deletants of P38 and P27. We also describe a cellular assay for antagonism of PACAP’s action at PAC1 that faithfully recapitulates PACAP antagonism by P6-38 in cell-based assays [52] and as reported using P6-38 as an antagonist for PACAP actions *in vivo* [53–60].

## 4. Discussion

PACAP, acting through PAC1, is involved in pathogenesis and progression of several disease states, including chronic pain, migraine, and stress-related conditions such as depression and PTSD [1, 63, 68–74]. Therefore a search for both small-molecule and peptide-based PAC1 antagonists has been extensive, including both physical screening of compound libraries and B1 GPCR analogs, and chemiformatic screening based on receptor and compound structural parameters [20, 25, 26, 31, 32, 75, 76]. Four SMOLs have been described that inhibit PACAP-dependent signaling in cell based assays [20, 21, 25, 26, 64]. Two of these are also reported to be effective in inhibiting PACAP-dependent nociceptive responses and restraint-induced anxious behaviors [63, 64]. However, we were unable to find evidence for specific PACAP antagonism for any of these compounds using a non-neuroendocrine cell line expressing exogenous hPAC1hop. Upon further examination, it was revealed that the HEK293_CBS_hPAC1-based assay did not exhibit inhibition of PACAP’s action by the only well-validated and specific PAC1 peptide inhibitors, P6-38 and the maxadilan-based M65. This motivated us to seek a PAC1-expressing cell line in which these peptides could inhibit P27 and P38 signaling through PAC1. The SY5Y_CBS cell line, expressing the endogenous PAC1null variant, the most abundantly expressed isoform in human brain and peripheral nervous system [61, 77, 78], allowed comparison of the potencies of putative SMOL inhibitors with peptide inhibitors. Under these conditions, the four SMOLs currently proposed as specific PAC1 inhibitors were far less potent, and not specific, for PACAP inhibition. The fact that these compounds appear to be highly potent for inhibiting PACAP signaling in other assay systems, including CHOK1 cells, and in blockade of nominally PACAP-dependent physiological events, such as allodynia, raise questions about their actual mechanisms of action, and whether these can be distinguished by further analysis. Co-crystallization of these compounds with PAC1 and testing in additional assays including non-specific cAMP signaling controls, such as forskolin or PAC1 non-expressing cells, would be highly informative for further understanding of ligand-receptor interactions in cellular screening [25] assays and other orthogonal testing strategies for validating SMOL PAC1 antagonists in future. Dengler et al. have pointed out that allosteric modulators of the closely related secretin receptor, for example, show pronounced cell specificity when their effects are compared in cell lines with endogenous receptor expression and exogenous receptor-expressing cell lines used for screening [79]. Additional assessment with the many alternately spliced isoforms of PAC1 would also be informative.

Several mechanisms for the differential effects of P6-38 and M65 on inhibition of PACAP signaling can be considered. The PAC1 ICL3 receptor subtype involved (hip, hop, hiphop or null) seems the least likely factor. Although PAC1 receptor subtypes, in particular the hop cassette, have been implicated in differential coupling to G_q_ and for calcium-mobilizing effects of PACAP-PAC1 signaling, coupling to G_s_ seems to be a common property of all of the PAC1 isoforms. G_s_-coupling that leads to elevation of cAMP and/or CREB phosphorylation is the read-out for our cell-based assay, and that of most other investigators who have developed peptide-based antagonists or SMOLs [20, 21, 25, 26, 64]. Differential ECD isoform effects on P6-38 binding have not been reported, albeit PACAP agonist effects are reported to vary between full and short ECD forms of PAC1 [80]. The differential expression of RAMPs is a potential explanation for differential properties of PAC1 interaction with PACAP, maxadilan, P6-38, and M65. RAMPs, or receptor activity-modifying proteins, are known to be important in selectivity of the calcitonin and calcitonin-like receptors CALCR and CALCLR, which with co-expression of RAMPs 1, 2, or 3 form distinct receptors for CGRP, calcitonin, amylin, and adrenomedullin [81, 82]. RAMP interaction with PAC1 has been inferred through multiplex analysis [83], however, specific examples of RAMP-PAC1 interaction leading to altered ligand recognition have not been reported. It is noteworthy that HEK293 cells show weak native expression of RAMPs 1 and 2 and little or no expression of RAMP 3 [84], whereas RAMP2 mRNA expression is high in both NS-1 and SH- SY5Y cells (Wu et al., unpublished observations). Further attention to this issue is warranted in order to better predict the *in vivo* pharmacology of both SMOL and peptide-based putative PAC1 inhibitors [85]. Finally, receptor-G protein stoichiometry in cells endogenously expressing receptor, and its effects on the binding poses available to the largest fraction of receptor presented on the surface of the cells being used for screening, is a potential factor in PACAP-PAC1 pharmacology.

Peptide-based family B1 agonists and antagonists have achieved dramatic clinical results in the past decade [86]. The results reported here demonstrate proof-of-principle for further development of peptide-based PACAP antagonists using dynamic modeling based upon empirically-derived structural information [13]. Current modeling suggests that amidation may not be required for inhibition, and is not required for PACAP agonist action as determined empirically[52]. C-terminal modification of P6-30 to the des-amide would allow potential delivery of the peptide via viral vectors for translational experimentation, which is not possible for the amidated peptides unless delivered specifically to a neuronal prohormone processing pathway. P6-30 is a potential scaffold for C-terminal extension to enhance binding of this inhibitory peptide to neighboring positively charged residues within the radius of interaction predicted by MD analysis of binding of P6-38, and P6-30, to PAC1 at the ECD and ECL predicted contact points for initial PACAP high-affinity trap binding to PAC1.

Discrepancies with previous reports of small-molecule and peptide antagonism may be explained, in part, by the use of assays that, while reflecting competition with some modes of PACAP binding to its receptor, may not recapitulate PACAP-antagonist competition leading to biological outcomes reflected in final activation of adenylate cyclase in the native environment of neuronal or neuroendocrine cells. The clearest example of this was our finding that P6-30, as predicted from structural analysis and molecular modeling, retained an antagonist potency equivalent to that of P6-38, despite indications to the contrary from classical 125-I-PACAP:PAC1 competitive binding studies [14].

Importantly, previously proposed small molecular antagonists derived from *in silico* predictions based on the binding of PACAP to the ECD of PAC1 were without activity in our neuroendocrine/neuroblastoma-based in cell-based assays. In an assay measuring PACAP-induced neuritogenesis in NS-1 cells, P6-30 and P6-38 also functioned as antagonists, providing a second orthogonal and biologically relevant assay against which putative antagonists can be tested. Further investigation of the mechanisms of action of currently developed PAC1-directed SMOLs in non-endocrine cells, in which PAC1 is expressed exogenously, seems warranted. Differential results for PACAP antagonism when employed in cellular assays in different cellular backgrounds may ultimately reveal differential pathways for PAC1 activation by PACAP [87, 88] of material relevance in the design and translational deployment of allosterically- as well as orthosterically- directed PACAP antagonists.

**Table 1.** Summary of the MD systems composition.

| **System,**<br>**# of replicas**<br>**(total sampling)** | **# of**<br>**atoms** | **# of**<br>**POPC**<br>**molecules** | **# of water**<br>**molecules** | **# of NaCl**<br>**molecules** | **Box dimension**<br>**after**<br>**equilibration (Å)** |
| --- | --- | --- | --- | --- | --- |
| **PAC1:P6-27,**<br>**3 x 250 ns**<br>**(750 ns)** | 100414 | 182 | 23055 | 77 (Na+)<br>67 (Cl-) | 83.34558 x<br>83.89778 x<br>140.7654 |
| **PAC1:P6-27Lb,**<br>**3 x 250 ns**<br>**(750 ns)** | 100401 | 182 | 23052 | 78 (Na+)<br>67 (Cl-) | 83.58749 x<br>83.03023 x<br>140.1813 |
| **PAC1:P6-27Lbx1,**<br>**3 x 250 ns**<br>**(750 ns)** | 100393 | 182 | 23047 | 78 (Na+)<br>67 (Cl-) | 84.07697 x<br>84.52230 x<br>138.7528 |
| **PAC1:P6-27Lbx2,**<br>**3 x 250 ns**<br>**(750 ns)** | 100393 | 182 | 23047 | 78 (Na+)<br>67( Cl-) | 82.84015 x<br>83.27893 x<br>143.4416 |
| **PAC1:P6-38,**<br>**3 x 250 ns**<br>**(750 ns)** | 96456 | 182 | 21670 | 67 (Na+)<br>63 (Cl-) | 82.70021 x<br>83.1382 x<br>137.5490 |
| **PAC1:P6-38Lb,**<br>**3 x 250 ns**<br>**(750 ns)** | 100538 | 182 | 23027 | 72 (Na+)<br>67( Cl-) | 83.01241 x<br>83.45210 x<br>142.4725 |
| **PAC1:P6-38Lbx1,**<br>**3 x 250 ns**<br>**(750 ns)** | 100225 | 181 | 22965 | 72 (Na+)<br>67( Cl-) | 84.14147 x<br>84.58714 x<br>138.5041 |
| **PAC1:P6-30,**<br>**6 x 250 ns**<br>**(1500 ns)** | 96539 | 182 | 21749 | 71 (Na+)<br>63( Cl-) | 83.87174 x<br>84.31598 x<br>133.9410 |
| **PAC1:(R)-PA-8 (1:7)<sup>a</sup>,**<br>**4 x 1000 ns**<br>**(4000 ns)** | 96408 | 180 | 21809 | 78 (Na+)<br>63( Cl-) | 83.74889 x<br>84.19863 x<br>134.2448 |
| **PAC1:(S)-PA-8 (1:7)<sup>a</sup>,**<br>**4 x 1000 ns**<br>**(4000 ns)** | 96408 | 180 | 21809 | 78 (Na+)<br>63( Cl-) | 83.49495 x<br>83.94332 x<br>134.9745 |
| **PAC1:(R)-PA-9 (1:6)<sup>a</sup>,**<br>**4 x 1000 ns**<br>**(4000 ns)** | 96279 | 180 | 21809 | 78 (Na+)<br>63( Cl-) | 83.69285 x<br>84.14228 x<br>133.9037 |
| **PAC1:(S)-PA-9 (1:7)<sup>a</sup>,**<br>**4 x 1000 ns**<br>**(4000 ns)** | 96322 | 180 | 21809 | 78 (Na+)<br>63( Cl-) | 83.83135 x<br>84.28153 x<br>134.0329 |
| **VPAC1:(S)-PA-9 (1:9)<sup>a</sup>,**<br>**8 x 500 ns**<br>**(4000 ns)** | 100382 | 179 | 23287 | 67 (Na+)<br>67 ( Cl-) | 83.59527 x<br>83.92934 x<br>14.10215 |

a = stoichiometry between PAC1 and the SMOLs in the simulation box.

## Acknowledgements

This work was supported by National Institute of Mental Health MH002386, Intramural Research Program of the National Center for Advancing Translational Sciences, NIH and the

Centre for Health and Life Sciences. PMS is a Leadership Fellow of the Australian National Health and Medical Research Council of Australia (ID: 2025694).

## Notes

### Competing Interest Statement

The authors have declared no competing interest.

### Summary of Updates

Figure 6 has been revised to correct an error in Figure 6C that was present in the initial submission.

